# Monitoring the circadian clock in human blood using personalized machine learning

**DOI:** 10.1101/066126

**Authors:** Jacob J. Hughey

## Abstract

The circadian clock and the rhythms it produces are crucial for human health, but frequently perturbed by the modern environment. At the same time, circadian rhythms may influence the efficacy and toxicity of therapeutics and the metabolic response to food intake. Measuring the body’s response to treatments for circadian dysfunction, as well as optimizing the daily timing of treatments for other health conditions, requires a simple and accurate method for monitoring the circadian clock. Here we used a recently developed method called ZeitZeiger to predict circadian time (CT, time of day according to the circadian clock) from genome-wide gene expression in human blood. In cross-validation on 498 samples from 60 individuals across three publicly available datasets, ZeitZeiger predicted CT in single samples with a median absolute error of 2.1 h. The predictor trained on all 498 samples used 15 genes, only two of which are part of the core circadian clock. We then extended ZeitZeiger to make predictions for groups of samples, and developed a general framework to personalize predictions using samples from only the respective individual. Each of these strategies improved prediction of CT by ~20%. Our results are an important step towards precision circadian medicine.

## Introduction

Much of human physiology, from sleep to immune function, has a daily rhythm (1). Driving many of these rhythms is a cell-autonomous molecular oscillator called the circadian clock, which is present in nearly every tissue in the body (2). In animal models, disrupting the circadian system genetically or environmentally can have a wide range of phenotypic consequences (3–5). In humans, circadian dysfunction is linked to a number of health conditions, including cancer (6), major depressive disorder (7), and obesity (8). At least some of the circadian dysfunction in humans seems to be a result of multiple features of the modern environment, e.g., shift work and reduced exposure to sunlight (9, 10). Consequently, improving circadian function by photic, behavioral, or other means, which has been called chronomedicine, could greatly benefit human health (11).

At the same time, increasing evidence suggests that circadian rhythms influence the efficacy and toxicity of therapeutics (12, 13) as well as the metabolic consequences of food intake (14). For example, over half of the 100 best-selling drugs in the U.S. target a protein whose mRNA in mice shows a circadian rhythm in at least one organ (15). Using knowledge of the body’s circadian rhythms to optimize the timing of interventions has been called chronotherapy (16, 17).

Large-scale implementation of chronotherapy may be more difficult than first thought, however, as multiple lines of evidence suggest that at any given time of day, different individuals’ circadian rhythms are at different points in the cycle. For instance, circadian phase of entrainment (as measured by the Munich Chronotype Questionnaire) varies highly between individuals (18), as well as with age and between day-workers and shift-workers (19). Furthermore, recent work found that diurnal preference correlated with the circadian phase of clock gene expression in hair follicle cells (20). These observations imply that the optimal timing of a given intervention may vary from one person to another.

Thus, one critical component of both chronotherapy and chronomedicine, which together might be called precision circadian medicine, is a method to determine the state of a person’s circadian clock(s) in real-time. For chronotherapy, such a method would be an input; for chronomedicine, an output. One method for detecting the state of the clock in humans is to measure the sleep-wake rhythm, either by questionnaires, sleep logs, or actigraphy (21). Although measuring the sleep-wake rhythm is non-invasive and has led to valuable insights (22,23), sleep-wake is also influenced by non-circadian processes and seems to have a complex relationship with the various clocks throughout the body (24).

The other common method for assessing the state of the clock is to measure melatonin in plasma or saliva. In particular, the dim light melatonin onset (DLMO) is the gold standard for circadian phase (25-27). Determining DLMO, however, requires collecting many samples under controlled conditions over at least several hours, making it impractical for widespread use or for monitoring the circadian clock in real time. Furthermore, DLMO only reflects the phase of the central clock in the suprachiasmatic nucleus (which controls secretion of melatonin by the pineal gland), making it unable to report on clocks in other tissues. In an effort to address these limitations, Ueda et al. previously developed the molecular-timetable method and used it to estimate CT from metabolite levels in human blood (28-30). Because the study was based on only six individuals under highly controlled conditions, however, the extent to which the approach will generalize to larger populations and diverse conditions has remained unclear.

More recently, we developed a machine learning method called ZeitZeiger and used it to achieve state-of-the-art prediction accuracy of CT in multiple mouse organs (31). ZeitZeiger’s prediction accuracy is enabled by three important features: fitting periodic behavior using splines instead of sinusoids, focusing on the variation in the data associated with the periodic variable of interest (e.g., time of day), and using regularization to prevent overfitting.

Here we applied ZeitZeiger to three publicly available datasets of genome-wide circadian gene expression in human blood. We found that ZeitZeiger learned to use a small set of genes to accurately predict the CT of a single sample. This allowed ZeitZeiger to detect how circadian gene expression is affected by various perturbations to the light-dark and sleep-wake cycles. We then investigated two ways to improve prediction accuracy: first, by using groups of samples, and second, by combining the initial prediction with that from a personal predictor trained only on samples from the respective individual. Our results are an important step towards precision circadian medicine.

## Results

### Predicting circadian time of single samples in three datasets

We assembled three publicly available datasets of genome-wide gene expression in human blood (Table 1) (32–34). Each dataset consisted of ~8 control samples taken throughout the day for each individual. We merged and batch-corrected the gene expression measurements (35, 36) and standardized the time of day values (Materials and Methods).

**Table 1.** Datasets of circadian gene expression in human blood. Numbers of samples include only those from the control condition, which were used for the majority of the analysis. ‘Dim’ corresponds to <10 lux in GSE39445 and <5 lux in GSE48113.

| Dataset | Reference | Light-dark regimen | Subjects | Samples | Interval | Age (mean ± sd) | Female |
| --- | --- | --- | --- | --- | --- | --- | --- |
| GSE39445 | (32) | Constant dim<br>(following LD 16:8) | 24 | 221 | 3 h | 27.5 ± 4.3 y | 42% |
| GSE48113 | (33) | Dim:dark 14.7:9.3<br>(following LD 16:8) | 22 | 147 | 4 h | 26.3 ± 3.4 y | 50% |
| GSE56931 | (34) | LD 14:10 | 14 | 130 | 4 h | 29.7 ± 8.9 y | 57% |

Using the combined data, we then performed 10-fold cross-validation, in which ZeitZeiger learned to predict a sample’s CT based on its gene expression. We ran cross-validation with a range of values for ZeitZeiger’s two main parameters, sumabsv (which controls the amount of regularization) and nSPC (which controls how many sparse principal components, SPCs, are used for prediction). Samples from the same individual were always in the same fold.

We evaluated the results of cross-validation in terms of absolute error (absolute difference between predicted and observed CT; Fig. 1A). The median absolute error achieved by the optimal parameter was 2.1 h (interquartile range 2.8 h). The expected absolute error of a random predictor is 6 h. Similar to our experience predicting CT using gene expression in mice (31), prediction accuracy plateaued at sumabsv=2 and nSPC=2. Prediction accuracy was similar across the three datasets, although somewhat worse in samples from GSE56931 (Fig. 1B). On average, the predictors from cross-validation trained with sumabsv=2 and nSPC=2 were based on the expression of 15 genes (Fig. 1C). These results suggest that ZeitZeiger can use the expression of a small number of genes in human blood to accurately predict circadian time from a single sample.

**Fig. 1.**
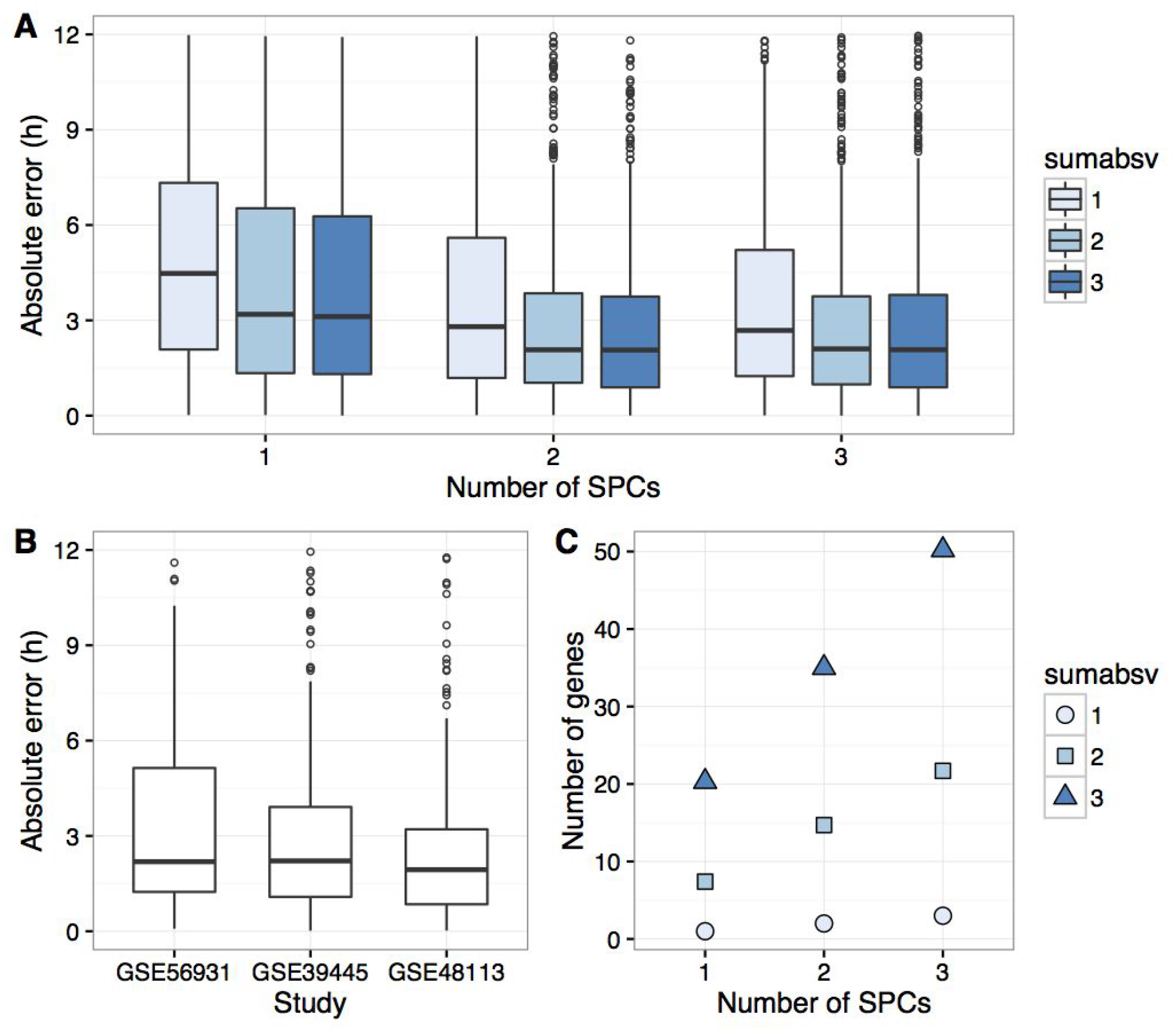
Using ZeitZeiger to predict circadian time across three datasets in 10-fold cross-validation. **(A)** Boxplots of absolute error for various values of sumabsv (regularization parameter) and nSPC (number of SPCs). **(B)** Boxplots of absolute error for each dataset at sumabsv=2 and nSPC=2. **(C)** Mean number of genes in the predictors from cross-validation for various values of sumabsv and nSPC.

Using the parameter values sumabsv=2 and nSPC=2, we trained a predictor on all control samples from the three datasets. The SPCs calculated by ZeitZeiger, each of which is a linear combination of genes, are designed to explain variation in gene expression associated with circadian time. The predictor’s two SPCs showed times of peak expression that were shifted from each other by ~6 h (Fig. 2A), similar to the multi-organ predictor of CT that we trained from gene expression in mice (31). Interestingly, however, the expression of SPC 1 as a function of CT was markedly non-sinusoidal. Moreover, of the 15 genes that formed the two SPCs (Fig. 2B), only two, *NR1D2* (*REV-ERBβ*) and *PER1*, are thought to be part of the core circadian clock. Consistent with this observation, the signal-to-noise ratio of circadian rhythmicity was generally lower for clock genes than for the 15 genes in the predictor (Fig. S1). When we allowed ZeitZeiger to predict CT using only core clock genes, absolute error on cross-validation increased by a median of 27% (P = 7 · 10^−6^ by paired Wilcoxon rank-sum test; Fig. S2), demonstrating ZeitZeiger’s ability to select the most informative genes.

**Fig. 2.**
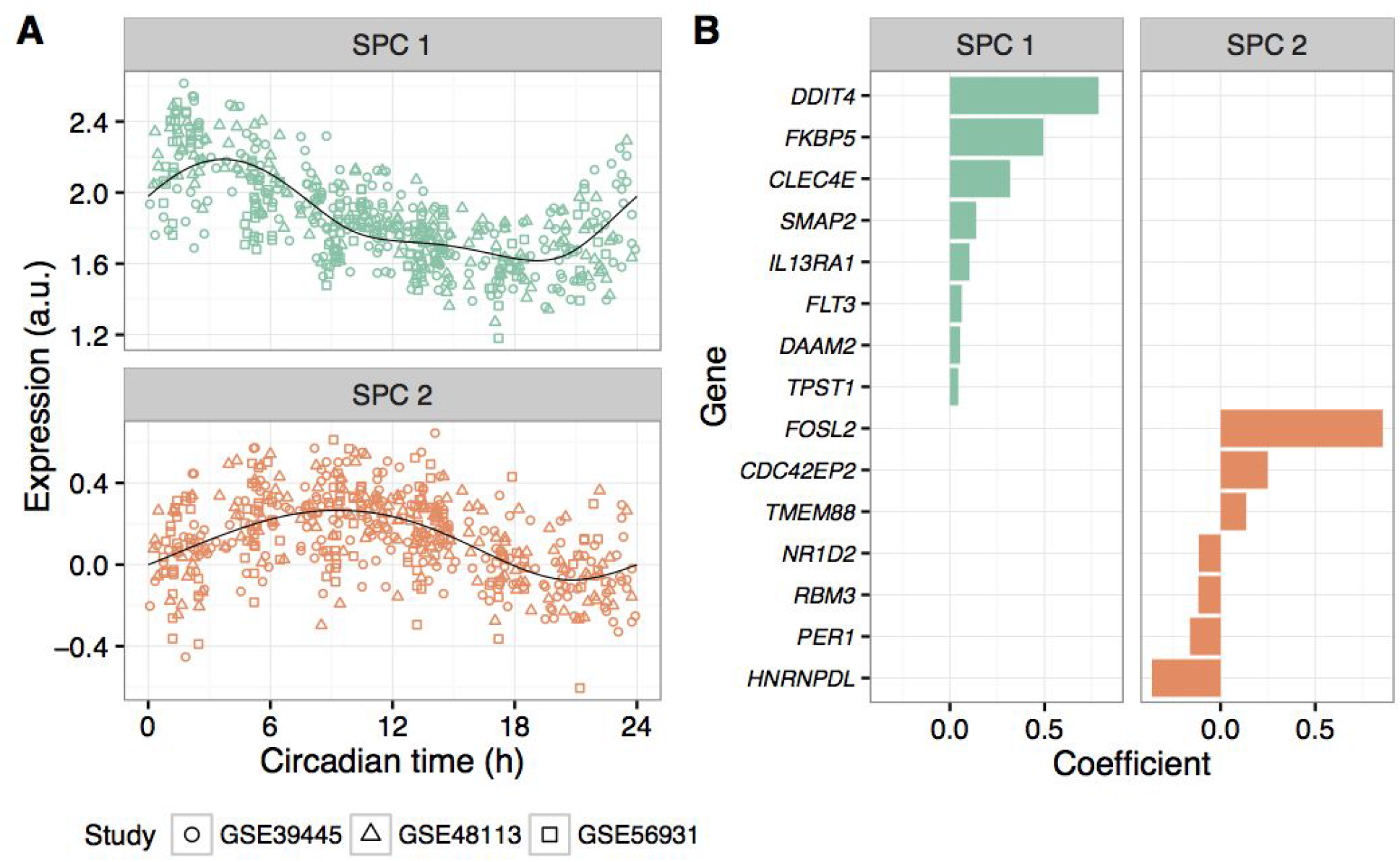
Properties of the predictor trained on all control samples across the three datasets (sumabsv=2, nSPC=2). **(A)** Expression of the two SPCs vs. circadian time. Each point is a sample. Black curves correspond to periodic smoothing splines fit by ZeitZeiger. **(B)** Genes and coefficients for the two SPCs. Genes are sorted by their respective coefficients. The expression of a given SPC in a given sample corresponds to the dot product of the coefficients for that SPC and the gene expression for that sample.

### Analyzing the effects of sleep perturbations

In addition to including samples from a control condition, each of the three datasets also included samples from a condition in which sleep (and the light-dark cycle) was perturbed. To investigate how the sleep perturbations would affect predictions of CT, we followed a leave-one-study-out strategy, in which we trained a predictor (sumabsv=2 and nSPC=2) on control samples from two datasets, then tested on all samples from the third dataset.

The sleep perturbations had several effects on predictions of CT (Fig. S3). First, six days of restricted sleep opportunity (6 h per night) worsened prediction accuracy by 16%, consistent with weaker circadian oscillations in gene expression (32). Second, a single night of complete sleep deprivation (38 h of continuous wakefulness with lights on (34)) appeared to induce a phase delay of 2.1 h. Finally, four 28-h days (forced-desynchrony protocol in GSE48113, which decouples the sleep-wake rhythm from the melatonin rhythm (33)) induced a phase delay of 2 h and increased the variability in prediction error by 42% (based on circular standard deviation).

We also used all the samples from GSE48113 (“in phase”, i.e., control, and “out of phase”) to train a predictor of time relative to DLMO, obtaining prediction accuracy similar to that obtained by predicting circadian time of control samples (Fig. S4A-C). The predictor trained on all samples from GSE48113 (sumabsv=2 and nSPC=2) included many of the same genes that were in the predictor trained on control samples from all three datasets (Fig. S4D-E). We therefore focused on the control samples for the remainder of our analysis.

### Predicting circadian time using multiple samples

Although ZeitZeiger was originally designed to predict the circadian time of a single sample, we wondered if prediction accuracy could be improved by using multiple samples from the same individual. We therefore extended ZeitZeiger to make predictions for groups of samples, where the time difference between each sample in the group is known (Materials and Methods). For each individual in the three datasets, we then constructed groups of two samples (taken either 8-9 h or 12 h apart) or three samples (taken over 12 h). We predicted the CT of each group in 10-fold cross-validation using sumabsv=2 and nSPC=2 (group information is only used during testing, not during training).

Compared to predictions based on a single sample, predictions based on two samples taken either 8-9 h apart were ~21% more accurate (0.43 h reduction in median absolute error; Fig. 3). Predictions based on two samples taken 12 h apart or on three samples showed a slight additional increase in accuracy (P > 0.3 by Wilcoxon rank-sum test). These results suggest that ZeitZeiger can use multiple relatively noisy samples to make better predictions.

**Fig. 3.**
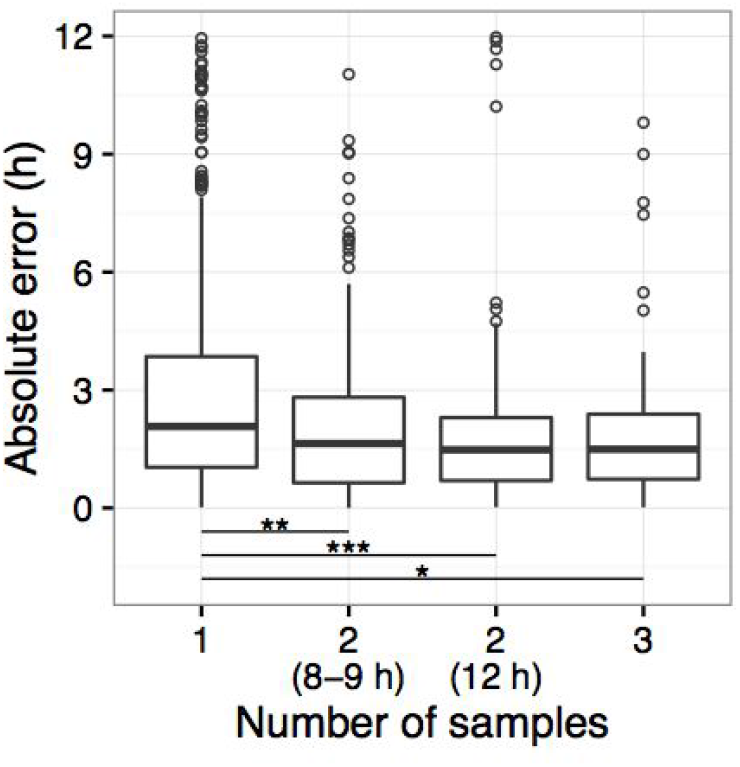
Predicting circadian time using various numbers of samples from the same individual. Boxplots of absolute error from 10-fold cross-validation. From left to right, the numbers of groups (one prediction per group) were 498, 198, 200, and 121. Levels of statistical significance (Wilcoxon rank-sum test): * P = 2.5 · 10^−4^, ** P = 1.9 · 10^−5^, *** P = 5.9 ·10^−7^. Other differences were not statistically significant.

### Personalizing predictions of circadian time

We previously found that ZeitZeiger can make accurate predictions even given small training sets with low time resolution (31). Because the three datasets included multiple samples per individual (median 8.5), we wondered if ZeitZeiger could learn to accurately predict CT using only the samples from a single individual. To test this, we performed leave-one-sample-out cross-validation for each individual (sumabsv=2 and nSPC=2). Unfortunately, the predictions from personal cross-validation were only slightly better than random and much worse than those from the original 10-fold cross-validation (Fig. S5).

Given that the merged dataset used throughout this paper included the expression of 17,477 genes, we suspected that ~8 samples per individual might not be enough to prevent ZeitZeiger from overfitting. We therefore devised a procedure for training personal predictors with “universal guidance”, which removes all features (i.e., genes) from the personal training set except for those selected by the “universal” predictor (i.e., the predictor trained on samples from multiple other individuals; Materials and Methods and Fig. 4A).

**Fig. 4.**
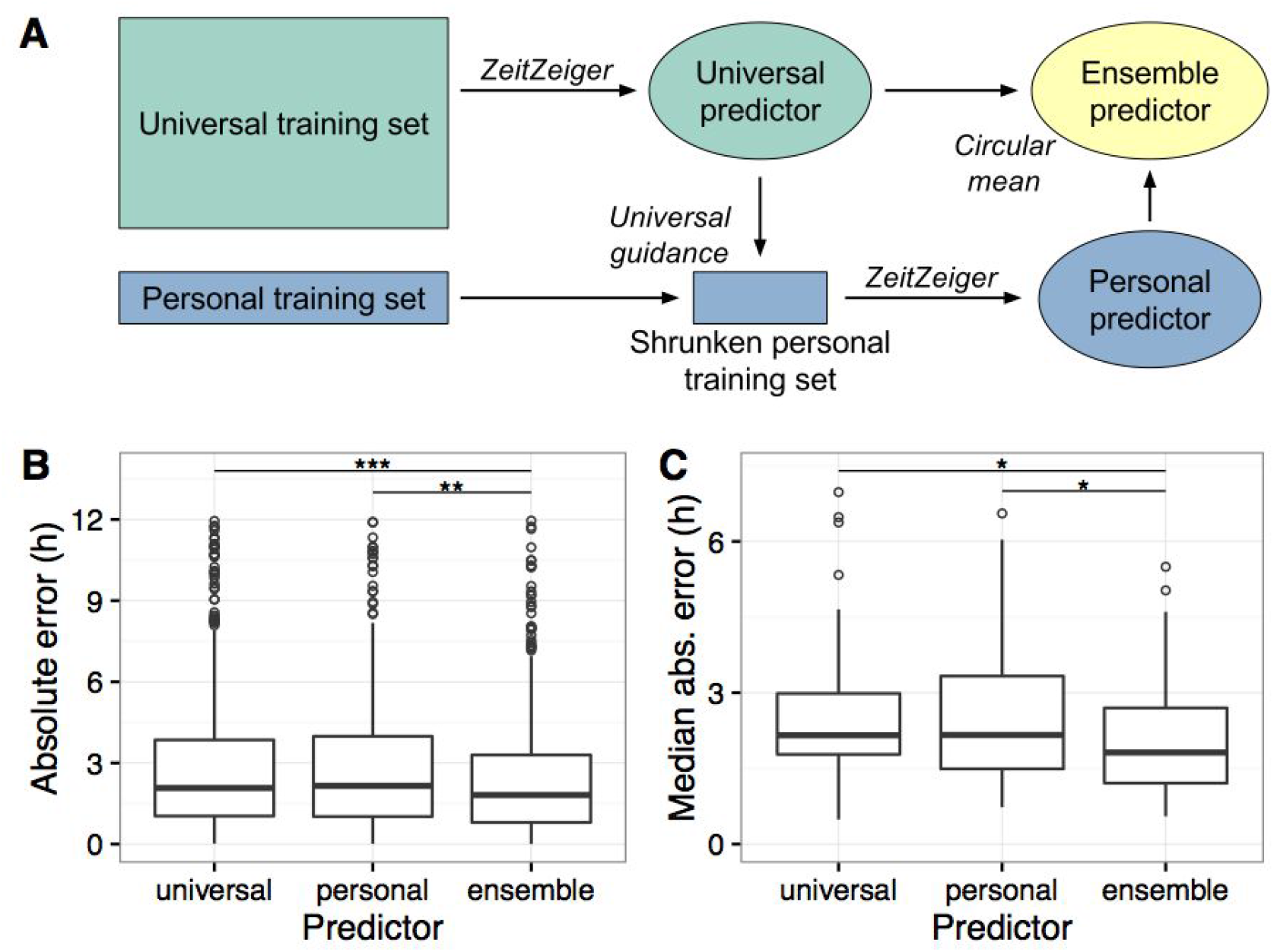
Personalized prediction of circadian time. All predictors were trained using sumabsv=2 and nSPC=2. Levels of statistical significance (paired Wilcoxon rank-sum test): * P < 10^−3^, ** P = 7.4 · 10^−5^, *** P = 5.9 · 10^−11^. **(A)** Schematic of procedure for training personal and ensemble predictors with universal guidance. For the training sets, the height represents observations and the width represents features (e.g., genes). Universal guidance refers to filtering for only those genes used by the universal predictor. **(B)** Boxplots of absolute error for universal (standard 10-fold cross-validation), personal (leave-one-sample-out cross-validation for each individual), and ensemble (circular mean of universal and personal) predictors. **(C)** Boxplots of median absolute error (by individual) for universal, personal, and ensemble predictors.

Using universal guidance, the personal predictors achieved similar accuracy on leave-one-sample-out cross-validation to the universal predictors on 10-fold cross-validation (Fig. 4B-C). We then combined the universal and personal predictors into an ensemble using the circular mean (Materials and Methods and Fig. 4A). Strikingly, the ensemble predictor was ~20% more accurate than either single predictor, both on a per-sample and per-individual basis (Fig. 4B-C and Fig. S6). The improvement in accuracy was robust for predictions based on at least seven personal training samples (Fig. S7). Applying this strategy of ensemble learning to groups of samples did not improve accuracy further, likely due to the small number of samples in the personal training sets (Fig. S8). Taken together, these results suggest that predictions based on a large training set from multiple individuals can be fine-tuned by predictions based on a carefully constructed training set from the individual of interest.

To further investigate the differences in circadian gene expression between individuals, we trained one predictor for each individual, using as universal guidance the 15 genes selected in the predictor trained on all samples. We found that the subset of genes selected by ZeitZeiger varied from one personal predictor to another (Fig. S9A). Furthermore, although the difference between peak times of SPC 1 and SPC 2 was largely consistent across individuals (Fig. S9B), the actual value of peak times was not (Fig. S9C). These results suggest that even the 15 “consensus” genes show meaningful interindividual variation in circadian expression.

## Discussion

Developing treatments that improve the function of or that account for the circadian system has the potential to improve multiple areas of human health. Realizing this potential, however, requires a robust method for monitoring an individual’s circadian rhythm. Here we developed a predictor of circadian time in human blood by applying machine learning to genome-wide gene expression. We demonstrated accurate prediction for single samples using a small set of genes, then developed two strategies that each improved accuracy by ~20%.

Both our strategies for improving prediction accuracy are based on the idea of an ensemble. Both rely on having multiple observations per individual. The first, using multiple samples with a known time difference between them, is effectively an ensemble of observations. The second, combining universal and personal predictors, fits the traditional definition of an ensemble in machine learning. In this case, the ensemble combines the relatively high-bias-low-variance universal predictor with the low-bias-high-variance (but not too high, thanks to universal guidance) personal predictor. This strategy for personalizing predictions is similar to an approach called customized training, which involves finding training observations that look similar to a given test observation (37). In the future, it may be possible to personalize predictions using samples from multiple individuals (e.g., individuals with similar chronotype or phase of entrainment).

Comparing our current results in human blood to our previous results in multiple mouse organs, two main differences emerge. First, our predictions here are less accurate, a consequence of circadian gene expression in humans being noisier (although here we analyzed only blood, we have observed similar levels of noise in human brain (38)). This increased noise is likely due to both genetic and environmental factors. The caveat is that some of the data from mice is based on tissue pooled from multiple animals, so a fair interspecies comparison of the variation in circadian gene expression in a given tissue has yet to be made.

Second, in contrast to the predictor we developed in mice, most of the genes in the human blood-based predictor are not thought to be part of the core circadian clock. This difference is likely due to the fact that the mouse predictor was trained on data from 12 organs, which discouraged ZeitZeiger from selecting genes whose circadian expression was tissue-specific and resulted in a strong enrichment for core clock genes. In this study, the dominant gene for SPC 1 was DDIT4 (REDD1), which encodes a protein that inhibits mTOR signaling as part of the response to cellular stress (39). The dominant gene for SPC 2 was FOSL2 (FRA2), which encodes a subunit of the AP-1 transcription factor and is therefore involved in numerous aspects of cell proliferation (40).

Because only two of the 15 genes in the predictor are part of the core clock and because circadian rhythms are normally aligned with the sleep-wake (and feeding) rhythm, it seems reasonable to wonder exactly which rhythm ZeitZeiger learned to detect. For three reasons, we believe the 15-gene ZeitZeiger predictor is primarily detecting of the progression of the clock. First, five of the ten core clock genes we analyzed showed a signal-to-noise ratio in the top 1.5% of all genes. Second, the control condition in two of the three datasets attempted to separate the clock from sleep, either by keeping subjects in constant conditions (GSE39445) or by allowing the clock to free-run (GSE48113). Third, we obtained similar results when training ZeitZeiger on control samples from all three datasets compared to training ZeitZeiger on all samples from the forced-desynchrony protocol of GSE48113. Further work is necessary to understand how the clock regulates the expression of the 13 non-core clock genes. Our results imply, however, that the forced-desynchrony protocol affects not just the central clock driving the melatonin rhythm, but also the peripheral clock in blood cells.

One limitation of this study is that, because not all of the datasets included individual-level information about DLMO, we had no direct measurement for the internal time of each individual’s circadian clock. Consequently, we trained ZeitZeiger to predict the externally measured time of day. Some of the inaccuracy of the predictions could therefore be due to interindividual variation in the alignment of external and internal time of day, i.e., the phase of entrainment. Such variation could explain why the personal predictors in the ensemble improved accuracy: they helped adjust for each individual’s circadian phase. Before our approach can be integrated into clinical trials for chronotherapy or chronomedicine, it will need to be validated in prospective studies. Such studies will likely involve testing ZeitZeiger and/or the 15-gene set alongside melatonin and actigraphy outside the laboratory setting.

Although here we have focused on genome-wide gene expression in blood, our methodology can be applied to any type of data from any tissue. We are therefore hopeful that in addition to its utility in chronotherapy and chronomedicine, ZeitZeiger will support efforts to study how circadian function in various tissues is altered in pathophysiological conditions and develop biomarkers for sleep- and circadian-related disorders (41). Furthermore, as the number of observations for each individual increases, e.g., in electronic medical records, our framework for personalizing predictions may prove useful in many areas of precision medicine.

## Materials and Methods

### Code and data availability

ZeitZeiger is available as an R package at https://github.com/iakeih/zeitzeiger. All data and code to reproduce this study are available at http://bit.ly/2a6M2HT (to be deposited at Dryad following publication).

### Processing time of day and other metadata

The nomenclature for time of day in chronobiology is complicated (42). Complicating our analysis even further, each of the three datasets used a different experimental design (in particular, a different light-dark regimen) and only one of the datasets included individual-level information for DLMO.

For both GSE48113 and GSE56931, the first samples were collected after subjects had been in the lab no more than a day. For GSE39445, the first samples were collected after subjects had been in the lab for 9 days, but for each subject, the midpoint of sleep opportunities in the lab coincided with the midpoint of sleep in that subject’s habitual sleep-wake schedule. In all three datasets then, the phase of each subject’s circadian clock should be based largely on the natural light-dark cycle. Therefore, we calculated the time of day for each sample in each dataset (e.g., 8:00 am) relative to sunrise time, using either the dates and geographic location provided by the authors (GSE56931) or the average sunrise time in the respective geographic location (GSE39445 and GSE48113). We refer to this adjusted time of day as “circadian time”.

For GSE48113, we calculated time relative to DLMO using the average DLMO for each condition (21:59 for “in phase”, 23:03 for “out of phase”), as provided in the original publication (33).

Unless otherwise specified, we used only the samples from the control condition in each dataset. This corresponded to “sleep extension” in GSE39445, “in phase with respect to melatonin” in GSE48113, and “baseline” in GSE56931.

### Processing the gene expression data

Gene expression from the three microarray datasets was processed using MetaPredict (36) (https://github.com/iakeih/metapredict), which maps probes to Entrez Gene IDs, performs intra-study normalization and log-transformation, and uses ComBat (35) to perform cross-study normalization. The merged data of control samples from all three datasets consisted of 17,477 genes measured in 498 samples.

### Using ZeitZeiger to predict circadian time

ZeitZeiger is a supervised learning method for periodic variables, i.e. variables that are continuous and bounded and for which the maximum value is equivalent to the minimum value. ZeitZeiger uses the training observations to learn a sparse representation of the variation associated with the periodic variable, then makes a prediction for a test observation using maximum likelihood (31).

Training a ZeitZeiger predictor involves the following steps: (1) fitting a periodic smoothing spline to the intensity of each feature (e.g., the expression of each gene) as a function of the periodic variable (43), (2) discretizing and scaling the spline fits, (3) using the discretized and scaled fits to calculate sparse principal components (SPCs; linear combinations of a small set of features) (44), and (4) fitting a periodic smoothing spline to the intensity of each SPC as a function of the periodic variable. Making predictions requires two steps: (1) projecting the test observation from feature-space to SPC-space and (2) using the spline fits of the SPCs from the training data to perform maximum likelihood estimation. The two main parameters of ZeitZeiger are sumabsv and nSPC. The former corresponds to the amount of *L*_1_ regularization used to calculate the SPCs, while the latter corresponds to the number of SPCs used for prediction. Other parameters of ZeitZeiger include the number of knots for spline fitting and the number of time-points for discretization. In this study, we always used three knots (which constrains the spline’s flexibility and makes it more resistant to noise) and 12 time-points.

Ten-fold cross-validation was performed such that all samples from a given individual were in the same fold. The folds were identical when predicting CT for groups of samples and when training universal predictors to provide universal guidance to the personal predictors in leave-one-sample-out cross-validation. Because only three datasets were available (two of which were from the same research group), we elected not to perform leave-one-study-out cross-validation or to have a separate group of validation samples from one or more of the datasets. Instead, we only performed 10-fold cross-validation across all controls samples from all three datasets. This means we may be underestimating generalization error (perhaps cancelling out the imperfect standardization of time of day), but also makes it simpler to use all the control samples when testing strategies for improving accuracy.

We did use a leave-one-study-out strategy when analyzing the effects of sleep-wake perturbations. For each dataset in turn, we trained a ZeitZeiger predictor (sumabsv=2 and nSPC=2) on only the control samples from two datasets, then tested on all samples (control and “treatment”) from the third. Thus, prediction accuracies from this analysis are not directly comparable to those from 10-fold cross-validation, but within this analysis, one can still compare results for control and treatment samples within each dataset.

The signal-to-noise ratio of circadian rhythmicity for gene *j* was calculated as

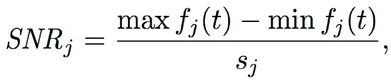

where *f_j_*(*t*) is the expression of gene *j* as a function of time *t*, and *s_j_* is the root mean squared error of the periodic spline fit.

### Providing universal guidance when training personal predictors

For each fold of 10-fold cross-validation (performed across individuals) and each sample of personal leave-one-sample-out cross-validation (performed on samples from a single individual), our procedure for universal guidance worked as follows (Fig. 4A). First, samples from the other nine folds (the universal training set) were used to train a “universal” predictor (the same predictor used for 10-fold cross-validation in Fig. 1). Next, the training samples from the current individual (i.e., the personal training set) were filtered to include only those genes used in the universal predictor, resulting in the shrunken personal training set. Thus, universal guidance here exploits the fact that ZeitZeiger performs feature selection. Finally, the shrunken personal training set was used to train the personal predictor.

### Extending ZeitZeiger for multiple samples and multiple predictors

When making a prediction for a single sample *x*, ZeitZeiger calculates the log-likelihood *L*(*t*|*x*), where *t* ∈ [0, 1) is the scaled periodic variable (e.g., time of day). The predicted time 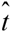 is then

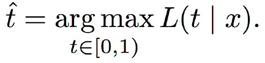

Now suppose we have a group of *n* samples, and for each sample, we have the measurements *x_i_* and a time difference τ*_i_*, which is the time of the *i* th sample relative to the time of a particular sample in the group. In the simplest case, *n* = 1 and τ = 0. For two samples taken anti-phase to each other, we could have τ_1_ = 0 and τ_2_ = 0.5. Now we can combine the log-likelihood for each sample and make one prediction for the entire group as follows:

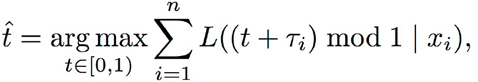

where the *mod* operator means that times less than 0 or greater than 1 “wrap around” to be between 0 and 1, and 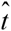 is the estimated time at which τ = 0.

Combining predictors can be done in two ways, the first of which works similarly to combining samples. Suppose we have a group of *m* predictors, where for a given sample *x*, *L_j_*(*t*|*x*) is the log-likelihood for predictor *j*. For the situation of one universal and one personal predictor, *m* = 2. The ensemble prediction is then

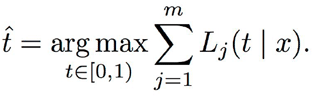

The second way to combine predictors is to use the circular mean. In this case, the ensemble prediction is

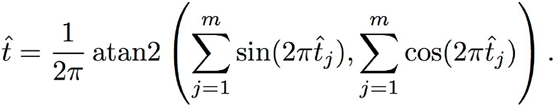

This second way is simpler and (on our data) provides a slightly larger improvement in accuracy, so all ensemble predictions in this study are based on the circular mean.

Our current implementations implicitly weight each sample or each predictor equally, but one could imagine incorporating explicit weights into any of these calculations, then learning the weights through an additional round of cross-validation.

## Acknowledgments

I thank Daniel Fabbri and members of Atul Butte’s lab for helpful comments. This work was supported by start-up funds from the Vanderbilt University School of Medicine.

## Author contributions

J.J.H. conceived and designed the study, performed the analysis, and wrote the manuscript. The 15-gene set has been disclosed for possible patent protection to the Vanderbilt Center for Technology Transfer and Commercialization by J.J.H.

## Supporting Information

**Fig. S1.**
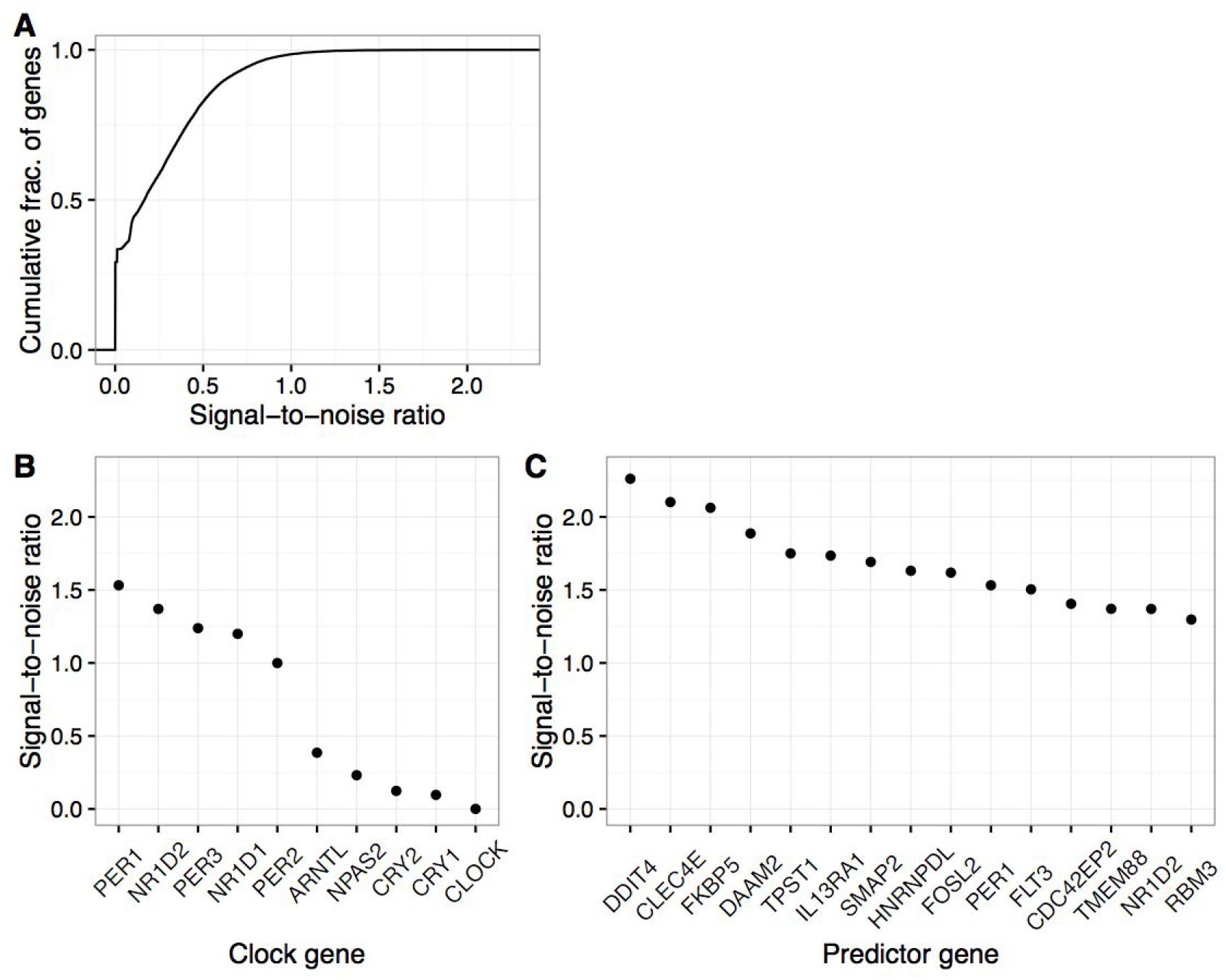
Strength of circadian rhythmicity, quantified as a signal-to-noise ratio (SNR), for gene expression in human blood. As in Fig. 2, data is from control samples from all three datasets. **(A)** Cumulative distribution function of SNR for all genes. **(B)** SNR for core clock genes. **(C)** SNR for genes in the ZeitZeiger predictor.

**Fig. S2.**
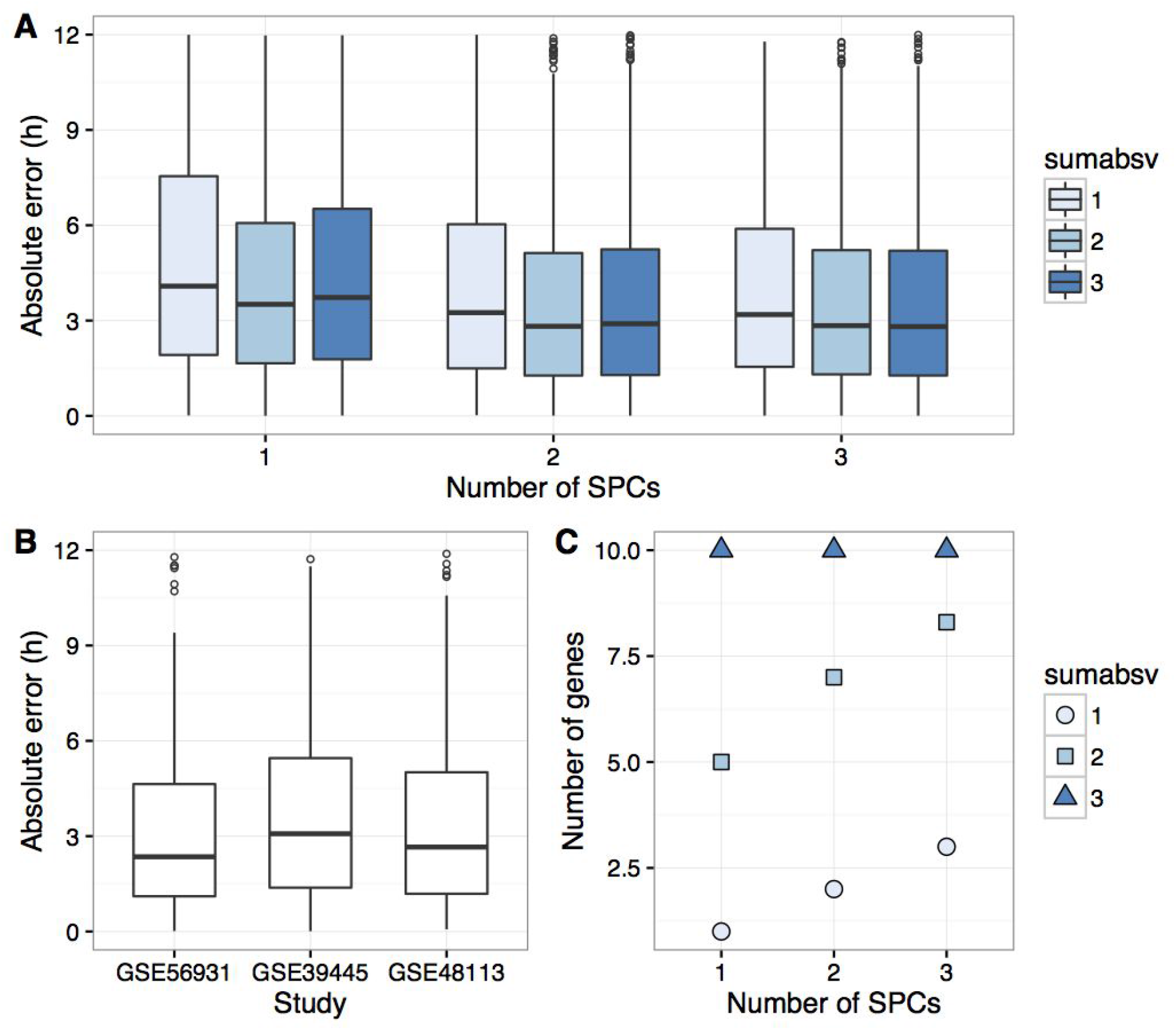
Ten-fold cross-validation to predict CT using only the core clock genes (otherwise identical to Fig. 1).

**Fig. S3.**
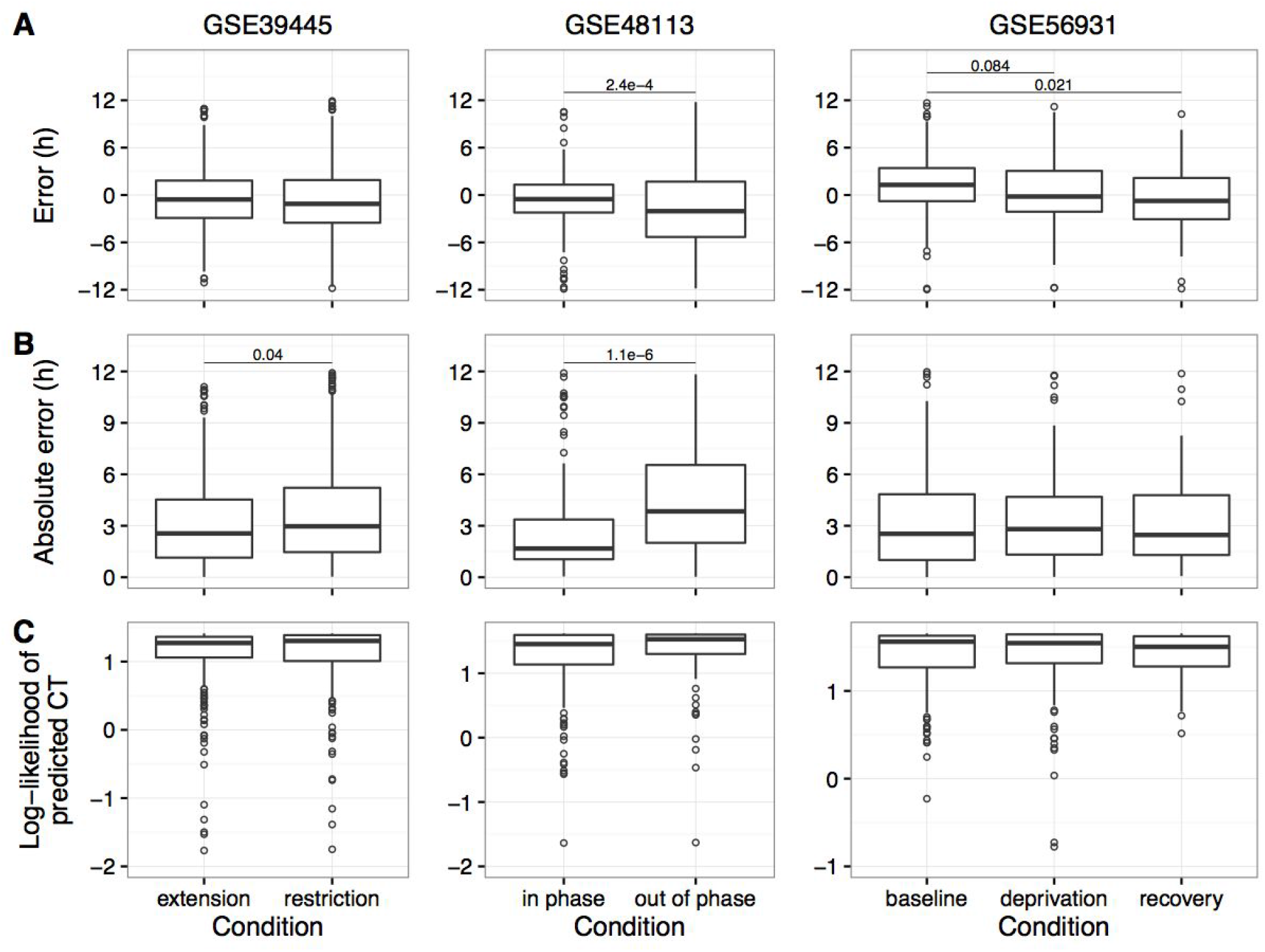
Applying ZeitZeiger to gene expression from various sleep perturbations. For each of the three datasets, a predictor was trained on control samples from the other two datasets, then tested on all samples from the dataset of interest. Boxplots of **(A)** error, **(B)** absolute error, and **(C)** log-likelihood of predicted circadian time for each condition in each dataset. Numbers above the boxplots indicate p-values less than 0.1 (circular ANOVA for error and Wilcoxon rank-sum test for absolute error). For ease of visualization, plots of absolute error do not show three outliers in GSE39445, one in GSE48113, and three in GSE56931. The left-most condition in each dataset is the control. In GSE39445, “restriction” refers to seven consecutive days of 6-h sleep opportunity per night. In GSE48113, “out of phase” refers to a forced-desynchrony protocol that decouples the sleep-wake rhythm from the central circadian clock. In GSE56931, “deprivation” refers to one full day of sleep deprivation, while “recovery” refers to a normal sleep opportunity the next night.

**Fig. S4.**
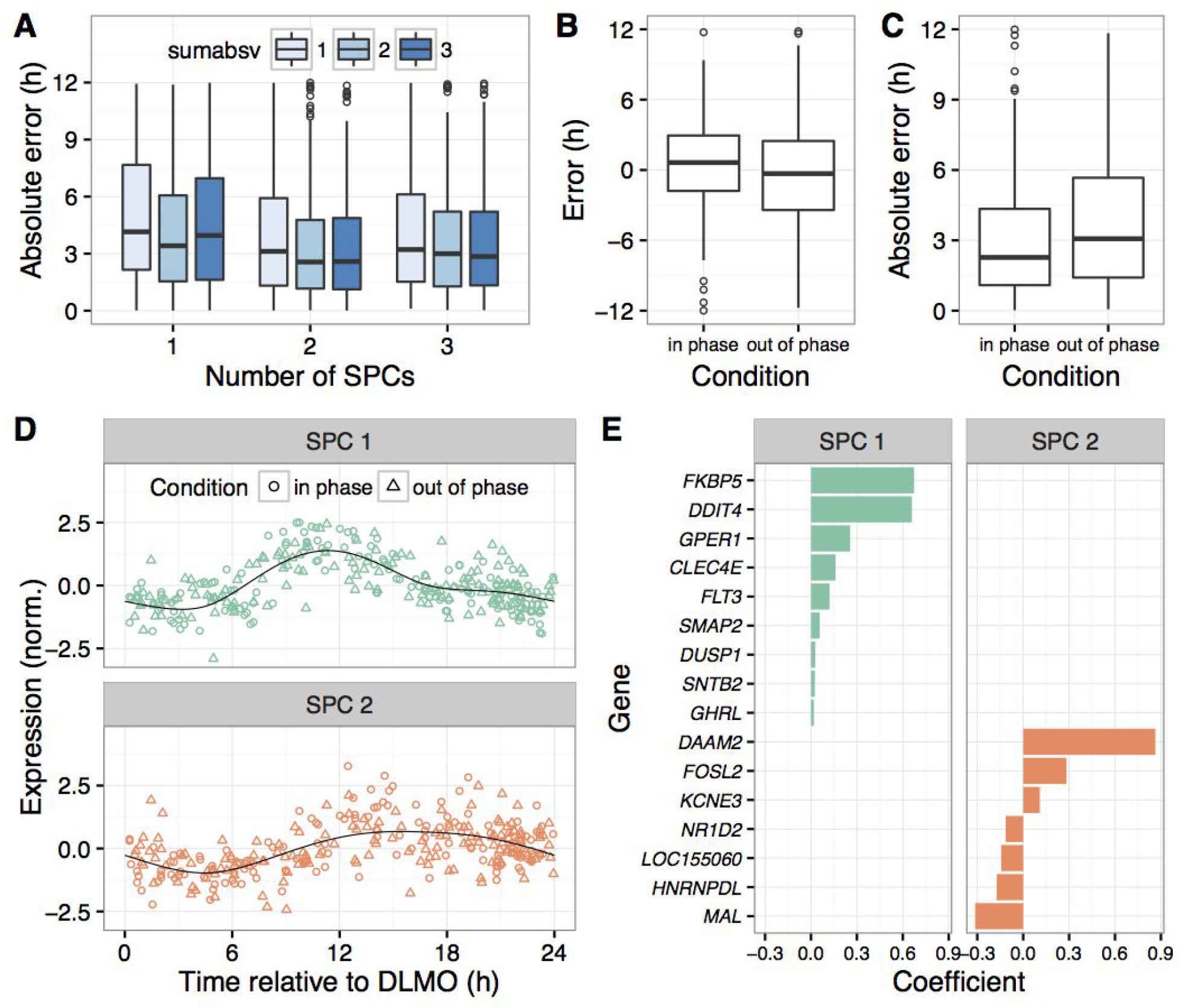
Applying ZeitZeiger to all samples (“in phase” and “out of phase”) from forced-desynchrony protocol of GSE48113. Instead of predicting circadian time, ZeitZeiger was trained to predict time relative to DLMO. **(A)** Boxplots of absolute error on 10-fold cross-validation for various parameter values. **(B)** Error and **(C)** absolute error on cross-validation (sumabsv=2 and nSPC=2) for each condition. **(D)** Expression of the two SPCs vs. time relative to DLMO (sumabsv=2). Each point is a sample. Black curves correspond to periodic smoothing splines fit by ZeitZeiger. **(E)** Genes and coefficients for the two SPCs. Genes are sorted by their respective coefficients.

**Fig. S5.**
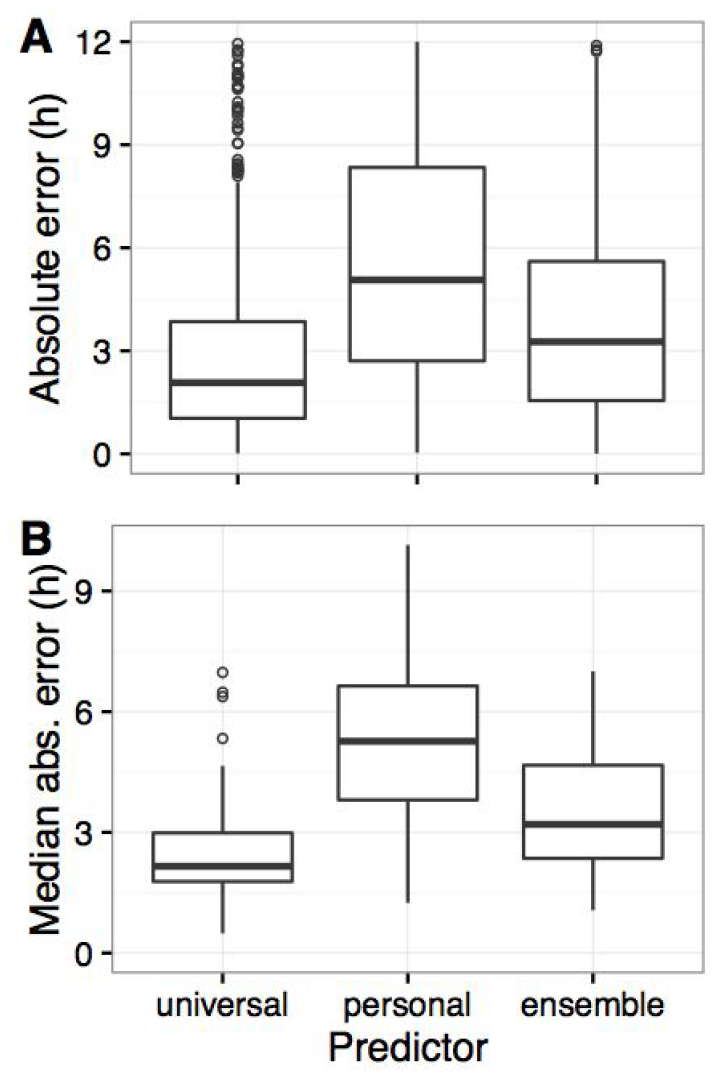
Boxplots of **(A)** absolute error and **(B)** median absolute error by individual for universal, personal, and ensemble predictors without universal guidance.

**Fig. S6.**
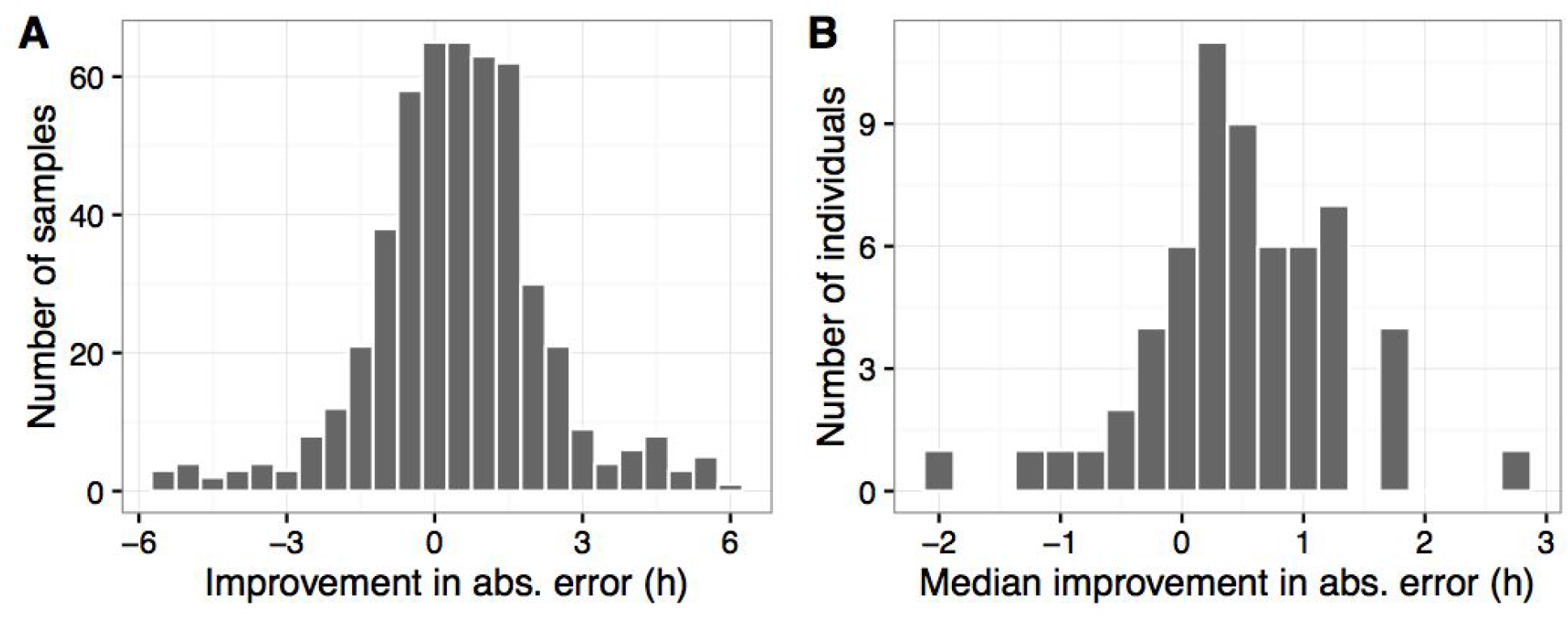
Histograms of **(A)** improvement in absolute error and **(B)** median improvement in absolute error by individual between the universal predictor and the ensemble predictor with universal guidance.

**Fig. S7.**
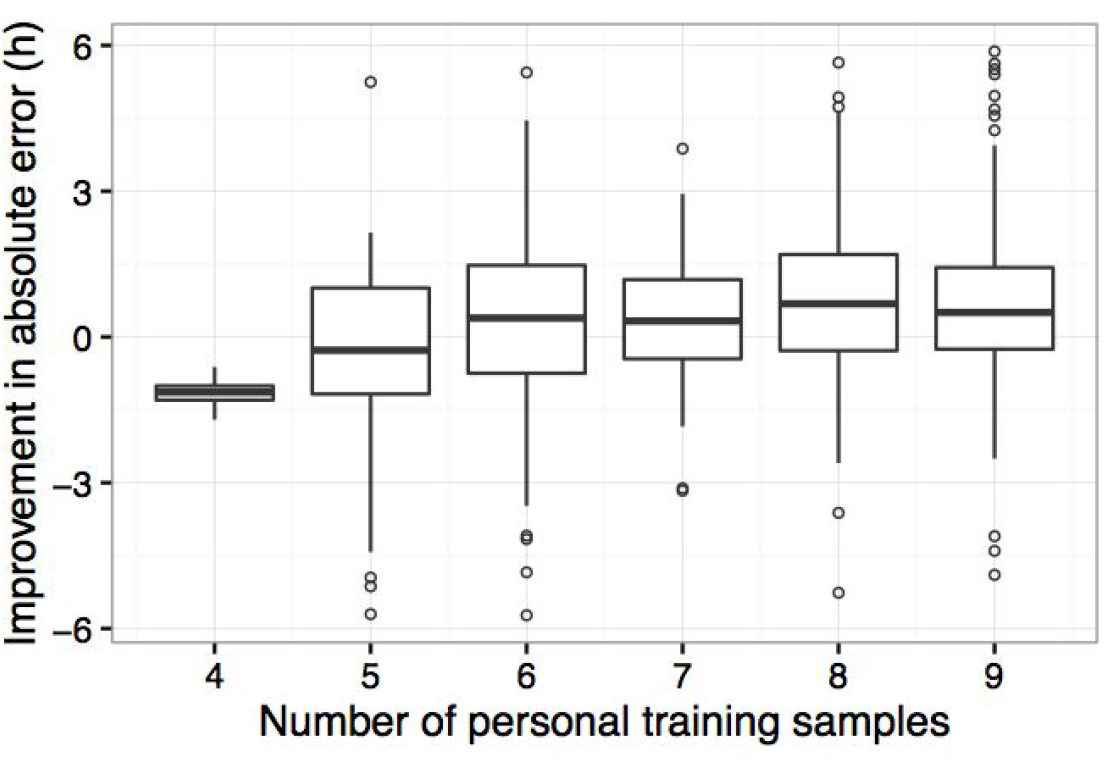
Boxplots of improvement in absolute error between the universal predictor and the ensemble predictor with universal guidance, as a function of number of personal training samples (because personal predictions were based on leave-one-out cross-validation, this is equal to the number of samples for the respective individual minus one).

**Fig. S8.**
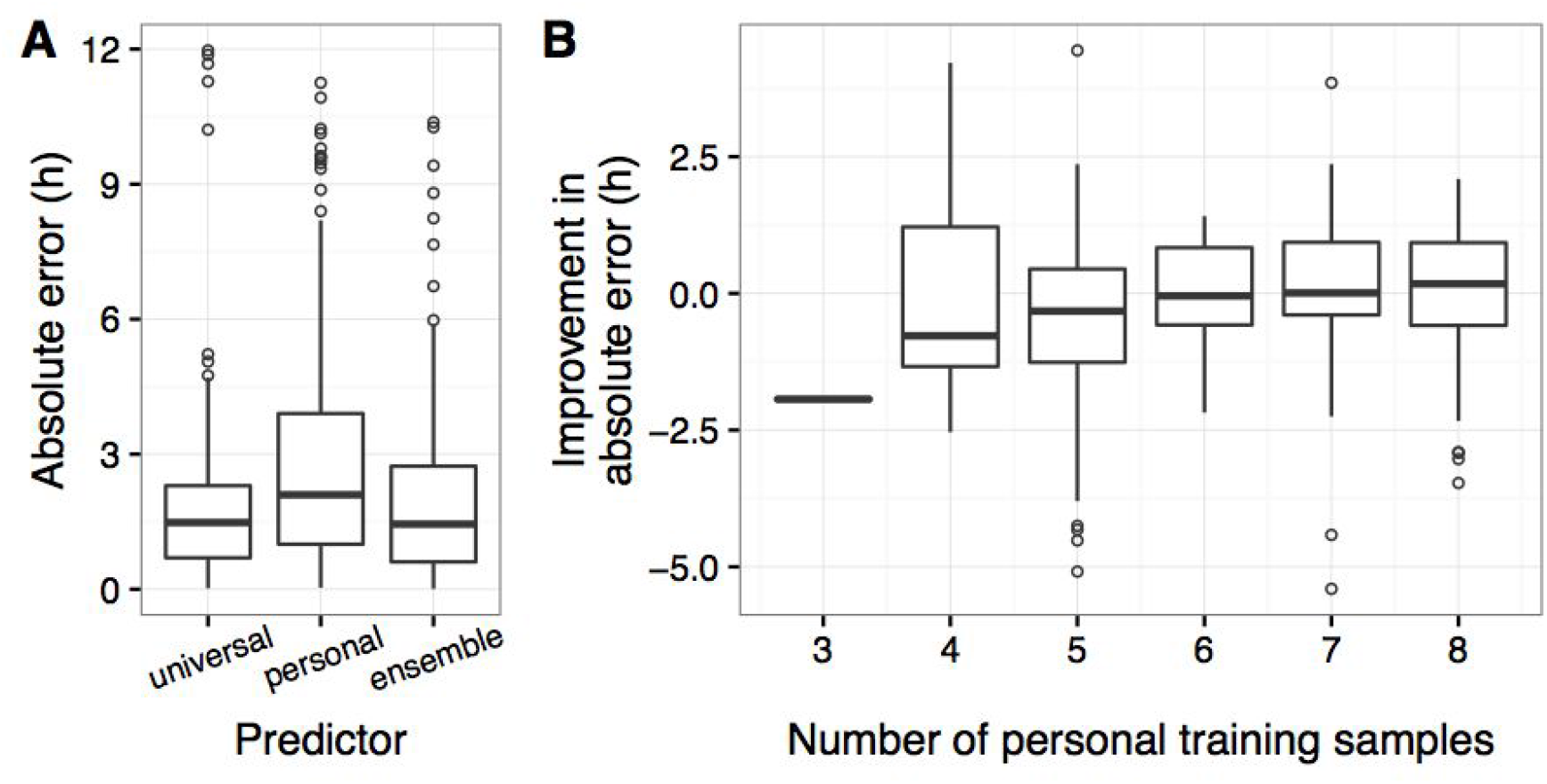
Personalized predictions with universal guidance applied to groups of samples. Each group consisted of two samples taken ~12 hours apart from the same individual. **(A)** Boxplots of absolute error for universal (standard 10-fold cross-validation, identical to Fig. 3), personal (leave-group-out cross-validation for each individual), and ensemble (circular mean of universal and personal) predictors. **(B)** Improvement in absolute error between universal predictor and ensemble predictor as a function of the number of personal training samples for that group (equal to the number of samples for that individual minus two).

**Fig. S9.**
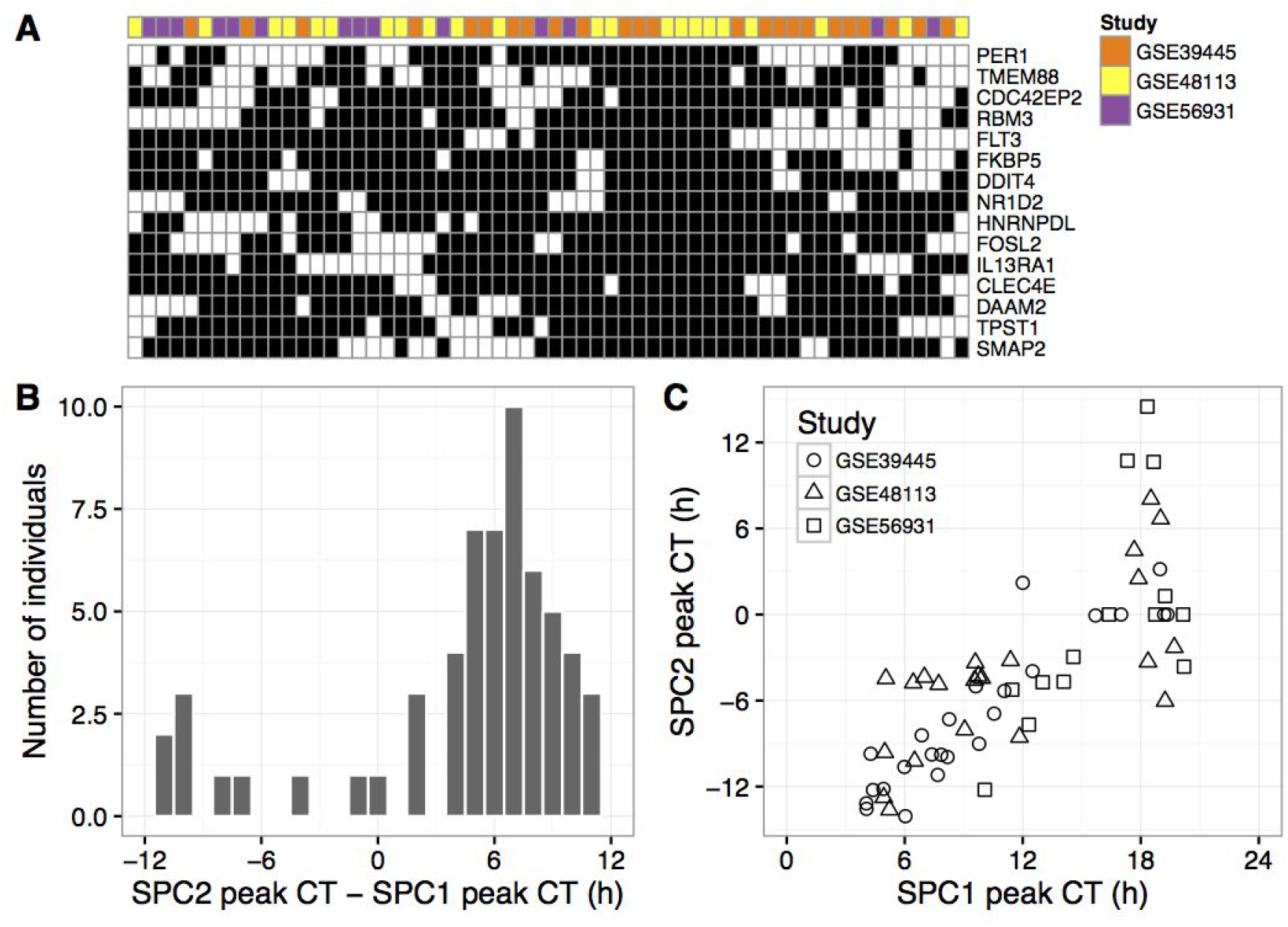
Genes and SPCs of the personal predictors trained with universal guidance. **(A)** Heatmap of genes present in personal predictors trained with universal guidance (using 15 genes present in predictor shown in Fig. 2). Rows correspond to genes and columns correspond to individuals. Black indicates the gene was present in the predictor for that individual. Rows and columns were sorted by hierarchical clustering. **(B)** Histogram of difference between peak times of SPC 1 and SPC 2. **(C)** Circadian times of peak expression for SPC 1 and SPC 2 in the personal predictors. Each point corresponds to one individual. For ease of visualization, some peak times for SPC 2 were shifted by 24 hours.

